# A two-stage ECG signal denoising method based on deep convolutional network

**DOI:** 10.1101/2020.03.27.012831

**Authors:** Lishen Qiu, Wenqiang Cai, Jie Yu, Jun Zhong, Yan Wang, Wanyue Li, Ying Chen, Lirong Wang

## Abstract

Electrocardiogram (ECG) is an effective and non-invasive indicator for the detection and prevention of arrhythmia. ECG signals are susceptible to noise contamination, which can lead to errors in ECG interpretation. Therefore, ECG pretreatment is important for accurate analysis. In this paper, a method of noise reduction based on deep learning is proposed. The method is divided into two stages, and two corresponding models are formed. In the first stage, a one-dimensional U-net model is designed for ECG signal denoising to eliminate noise as much as possible. The one-dimensional DR-net model in the second stage is used to reconstruct the ECG signal and to correct the waveform distortion caused by noise removal in the first stage. In this paper, the U-net and the DR-net are constructed by the convolution method to achieve end-to-end mapping from noisy ECG signals to clean ECG signals. The ECG data used in this paper are from CPSC2018, and the noise signal is from MIT-BIH Noise Stress Test Database (NSTDB). In the experiment, the improvement in the signal-to-noise ratio *SNR*_*imp*_, the root mean square error decrease *RMSE*_*de*_, and the correlation coefficient *P*, are used to evaluate the performance of the network. This two-stage method is compared with FCN and U-net alone. The experimental results show that the two-stage noise reduction method can eliminate complex noise in the ECG signal while retaining the characteristic shape of the ECG signal. According to the results, we believe that the proposed method has a good application prospect in clinical practice.

## I. Introduction

According to the World Health Organization (WHO) [1], cardiovascular disease is the leading cause of death in the world. The increasing burden of cardiovascular disease has become a major public health problem. Electrocardiogram (ECG) is a method widely used in the field of cardiology to analyze the heart condition of patients [2]. Compared with other methods, it has the advantages of being effective, non-invasive, and low-cost. ECG signals are measured by surface electrodes placed on the patient’s skin, but they are often contaminated by various noises, such as muscle artifact (MA), electrode motion (EM), and baseline wander (BW). MA can obscure details in ECG data and weaken certain characteristics of heart disease. ST-segment deviation from baseline due to EM or BW can be misdiagnosed as myocardial infarction, coronary artery insufficiency, or other diseases. These noise signals may affect the subsequent diagnosis of the ECG [3], so getting rid of this noise is the first step in ensuring that heart disease is correctly diagnosed.

Many researchers have contributed a wealth of algorithms to the development of ECG noise reduction technology. These algorithms mainly include infinite impulse response (IIR) [4], finite impulse response (FIR) [5,6], adaptive filter [7,8,9], discrete wavelet transform [10,11,12], Principal Component Analysis (PCA) [13], Independent component analysis (ICA) [14], and empirical mode decomposition (EMD) [15,16].

The methods based on IIR, FIR, and adaptive filters can remove noise outside the ECG frequency range. However, the methods will fail if the frequency range of the noise overlaps with the ECG, and the Gibbs effect is prone to occur when the order of the filter is not set properly. The wavelet transform-based method is widely used in ECG denoising because it can characterize the time-frequency domain information of the signal. However, it is necessary to select a suitable wavelet basis function as well as a suitable number of decomposition layers according to the signal to be analyzed, which lacks a certain degree of self-adaptation. In addition, the characteristics of the residual noise are complex, and its distribution in the frequency domain is unknown, so it is difficult to remove using a wavelet method. Based on the methods of PCA and ICA, the derived mapping model is very sensitive to small changes in signal or noise. The EMD-based method will discard part of the IMF in the reconstruction, causing a certain loss of information. Moreover, the EMD algorithm has problems such as modal aliasing and endpoint effects, and the algorithm operation takes too long. In general, there are two disadvantages of the aforementioned algorithms. One is that it cannot remove noise in a form close to the waveform, and the other is that it lacks sufficient adaptability.

Recently, denoising algorithms based on denoising autoencoders (DAE) [17] [18] have been shown to have better performance than traditional denoising algorithms. However, the biggest problem of DAE algorithms is that they destroy ECG characteristic waveforms and cannot retain valid detailed information. The above analysis shows that there is still room for further improvement of existing ECG denoising methods.

To solve the above problems, we propose a new two-stage denoising method. In the first stage, we propose a convolutional network based on an improved one-dimensional U-net. The main role of U-net is to remove noise as much as possible. After the first stage, baseline wander and high-frequency noise will be largely eliminated, but it is inevitable that part of the detailed waveform of the ECG will also be discarded. Therefore, in the second stage, we propose a new detailed waveform recovery network called DR-net. Through a special network structure, the network can largely recover the detailed information discarded by the U-net in the first stage. With this two-stage method, complete denoising of the ECG is achieved.

To the best of our knowledge, our work is the first to propose the application of one-dimensional U-net to ECG signal denoising. It is also the first to propose a two-stage method of denoising and then recovering, and to design a DR-net for recovering characteristic waveforms. The innovation point of this article is how to improve one-dimensional U-net and design one-dimensional DR-net.

The rest of this article is organized as follows: in Section 2, we introduce the experimental data we used. In Section 3, we introduce the evaluation indicators used in this article. The details of the proposed two-stage method based on U-net and DR-net are described in Section 4, Section 5 describes the experimental results obtained, and Section 6 summarizes this work.

## II. Data Description

How to obtain clean ECG data is the core of the problem of using deep learning to denoise. Most original papers use data manually selected or randomly intercepted from the MIT-BIH Arrhythmia Database for training and testing, but there are four problems with this:

1. MIT-BIH data has only two leads, 48 groups, and lacks abnormal ECG data. It is difficult to fully evaluate the impact of denoising algorithms on waveform characteristics.
2. The MIT-BIH data itself has a considerable degree of noise signals.
3. In the case of a small amount of data, the ECG of the same patient will be intercepted repeatedly in multiple groups, even after random disruption, and data leakage will inevitably occur.
4. In some papers, the signal-to-noise ratio (SNR) is fixed at a certain number of decibels, which is convenient for comparison, but it is difficult to represent the generalization performance of the proposed method.

The ground-truth ECG used in this article was derived from ICBEB 2018 [19], and 1,379 high-quality single-lead signals were manually selected from the competition dataset by trained volunteers. These data came from different, random leads in the standard 12-lead ECG. The original sampling frequency was 500 Hz. In order to unify the sampling rate with the noise signal, it was resampled to 360 Hz. In the experiment, the length of each dataset was unified to 3,600 sampling points (10 s). In addition to normal situations, the data also included abnormal conditions (such as ventricular premature beats and right branch obstruction). Hence, the ECG signals we collected and screened were rich in variety and close to the actual clinical situation. We randomly divided 1,379 signals into a signal training set (965), a signal verification set (276), and a signal test set (138) at a ratio of approximately 7:2:1.

The noise used in this paper was selected from the MIT-BIH Noise Stress Test Database (NSTDB) [20] [21]. The database included three common types of noise: MA, EM, and BW. Each noise source consisted of two columns, with a length of 650,000 points and a sampling frequency of 360 Hz. To prevent data leakage, we divided the noise into three non-overlapping parts: the noise training set (70%), the noise verification set (20%), and the noise test set (10%). We randomly set the starting point of the signal when intercepting noise, and the length of each intercept was 3,600 sampling points (10 s). After interception, the noise were superimposed on the signal training set, signal verification set, and signal test set. The noise components in the three sets were similar but not exactly the same. We believe that this setting is similar to the real situation and can inform the system of similar noise signals when building a noise reduction system. In order to better verify the noise reduction effect and better simulate the ECG signals of patients under real conditions, we divided the experiment into three groups: Group1, Group2, and Group3. Each group of experiments contained MA, EM, and BW when generating noisy signals. However, different noise ratios were maintained in each group of experiments to ensure that each kind of noise was dominant in one of the groups. In the three sets of experiments, the generation of noisy signals was carried out as follows:

noise-convolved ECG1 = 0.6 × MA + 0.2 × EM + 0.2 × BW + ground-truth ECG (Group1)

noise-convolved ECG2 = 0.2 × MA + 0.6 × EM + 0.2 × BW + ground-truth ECG (Group2)

noise-convolved ECG3 = 0.2 × MA + 0.2 × EM + 0.6 × BW + ground-truth ECG (Group3).

Figure 1 shows the different forms of the same group of signals in Group1, Group2, and Group3. MA is the dominant type of noise in Group1, EM is dominant in Group2, and BW is dominant in Group3. Such experimental design is conducive to comparing the performance of the algorithm when different noises are dominant. In the end, each set of noisy ECG datasets contained a total of 965 segments for training, 276 segments for verification, and 138 segments for testing. Figure 1(a) displays the ground-truth ECG.

**Fig. 1.**
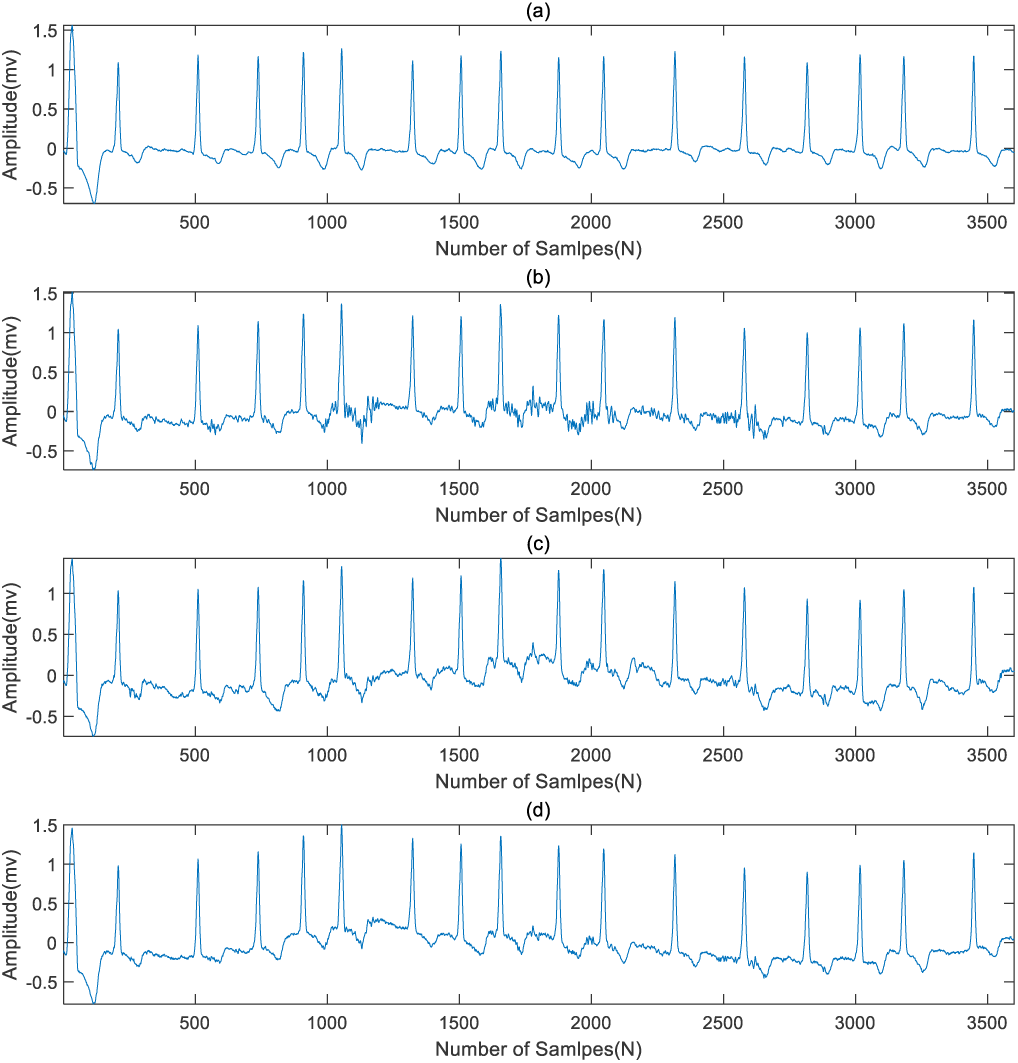
(a) Ground-truth ECG. (b) Noise-convolved ECG in Group1. (c) Noise-convolved ECG in Group2. (d) Noise-convolved ECG in Group3.

## III. Evaluation Indicators

In this study, three indicators, root mean square error (RMSE) decrease *RMSE*_*de*_ SNR increasing *SNR*_*imp*_, and Pearson correlation coefficient *P*, were used to evaluate the effect of denoising.

*RMSE*_*de*_ indicates the difference between the mean square error before noise reduction *RMSE*_*in*_ and the mean square error after noise reduction *RMSE*_*out*_. A larger *RMSE*_*de*_ indicates better noise reduction performance. *RMSE*_*de*_, *RMSE*_*in*_, and *RMSE*_*out*_ are defined as follows:

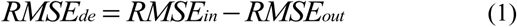

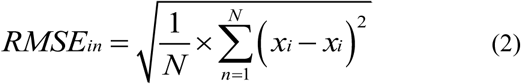

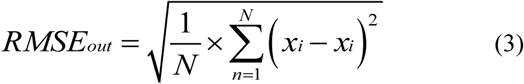

where *x*_*i*_ is the value of sampling point i in the ground-truth ECG, 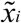 is the value of sampling point i in the noise-convolved ECG, and 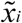 is the value of sampling point i in the ECG signal after denoising.

*SNR*_*imp*_ represents the difference between *SNR*_*out*_ and *SNR*_*in*_ of the input signal. A larger value indicates better noise reduction performance. *SNR*_*imp*_, *SNR*_*out*_, and *SNR*_*in*_ are described by the following expressions:

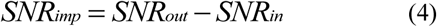

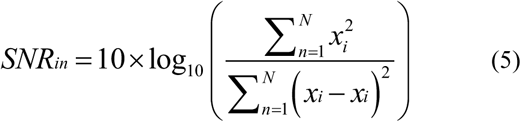

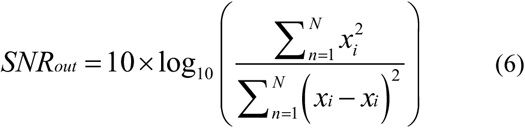

The QRS complex reflects the depolarization of the ventricle, and the T wave reflects the repolarization of the ventricle. These two waveforms play an important role in the diagnosis of ECG. We randomly selected 56 sets of signals in the test set, marking a total of 685 heartbeat QRS complex start points, end points, and T wave end points, and extracted the QRS complex and ST-T band. The relationship between ground-truth ECG and denoised ECG in terms of the QRS complex and ST-T band was evaluated by the Pearson correlation coefficient *P*.

*P* is described as

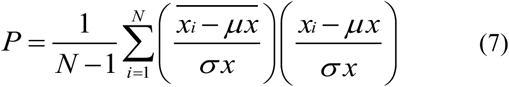

where *x*_*i*_ is the ground-truth ECG, 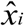 is the denoised ECG, *μx* and *σx* are the mean and standard deviation of *x*, respectively, and 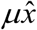 and 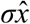 are the mean and standard deviation of 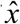, respectively.

## IV. Method

We designed a two-stage denoising method for ECG. The two stages correspond to two models. In the first stage, we designed an improved one-dimensional U-net model to denoise ECG signals. In the second stage, we designed a one-dimensional DR-net model and used this model to compensate for the characteristic waveform distortion and detail loss caused by the first stage. The overall flowchart of the algorithm is shown in Fig. 2.

**Fig. 2.**
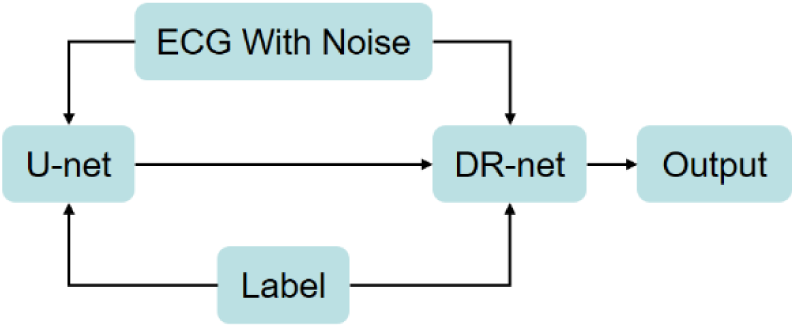
General flowchart of the proposed ECG denoising method.

U-net has proved its superior performance in image processing [22]. In [23], U-net was applied to one-dimensional ECG signals for the first time to realize R-wave localization and diagnosis of arrhythmia beats. We redesigned and improved the one-dimensional U-net network and applied it to the first stage of ECG filtering.

Our redesigned one-dimensional U-net model is shown in Fig. 3. Unlike the original model, there are three compression stages. The multiples of the three downsamplings are 1/10, 1/5, and 1/2, and the multiples of the three upsamplings are 2, 5, and 10. There are four sizes of convolution kernels, and the sizes from outside to inside are 30, 20, 10, and 5. According to [24], a large initial convolution kernel has a great effect on removing baseline drift. As the network deepens, the size of the convolution kernel decreases, which facilitates the network’s observation of details. On the left side of U-net, the encoding part, the convolution kernel gradually decreases. At the bottom of U-net, a context comparison mechanism [25] is added, as shown in the lower left corner in Fig. 3. This approach can increase the model’s attention to detail. On the right side of U-net, which is the decoding part, the convolution kernel increases in order. It is also worth noting that a skip connection from low-dimensional to high-dimensional is added to the model, and the skip connection is directly added to the corresponding convolution layer. This operation can alleviate information loss caused by upsampling and improve the denoising ability of the network. In the entire one-dimensional U-net network, a BatchNormalization (BN) layer is added to each hidden layer and passes through the activation function Relu.

**Fig. 3.**
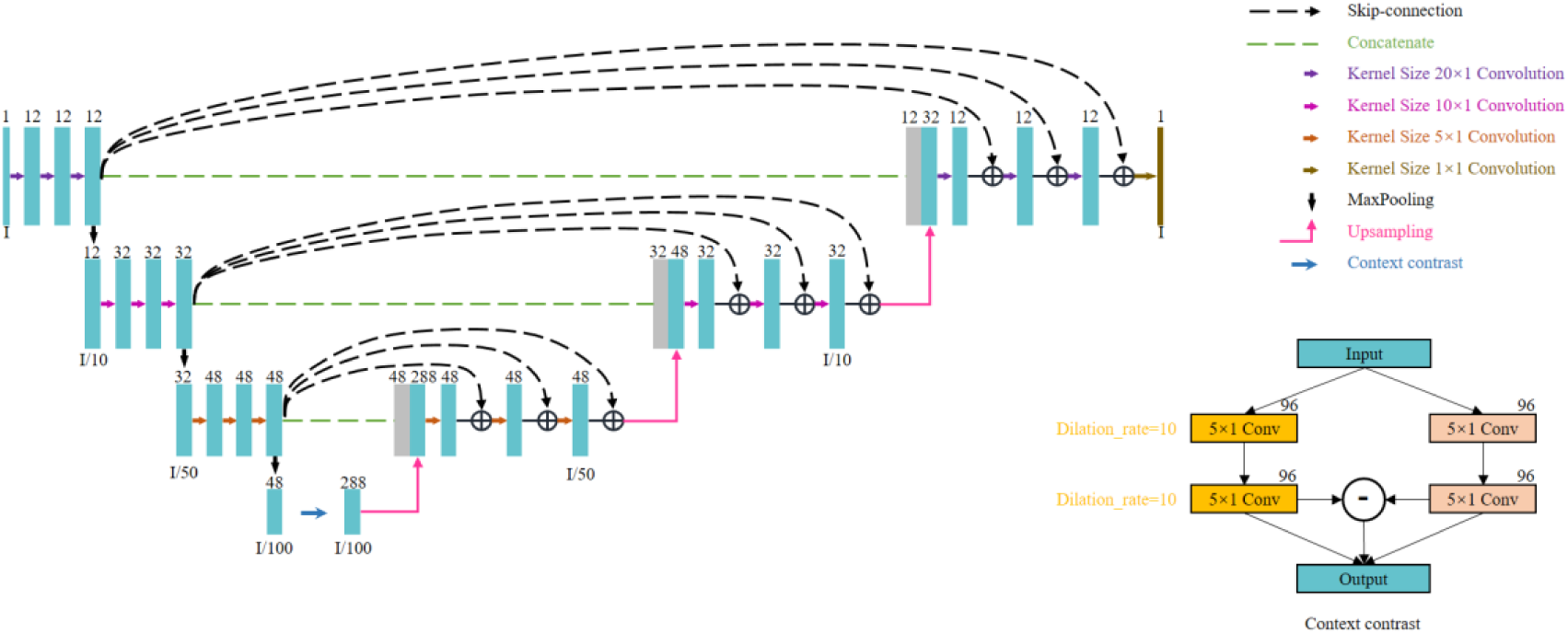
Improved one-dimensional U-net model structure.

After the first stage of denoising, the noise in the ECG signal will be eliminated, but the waveform of the ECG signal will inevitably be distorted to some extent. Especially in high-quality ECG, such changes are more obvious, as shown in Fig. 7. This distortion is largely caused by the downsampling layer in the network. The larger the multiple of the downsampling layer, the more obvious the distortion. However, if you reduce or cancel the downsampling layer, you cannot get good denoising results.

**Fig. 4.**
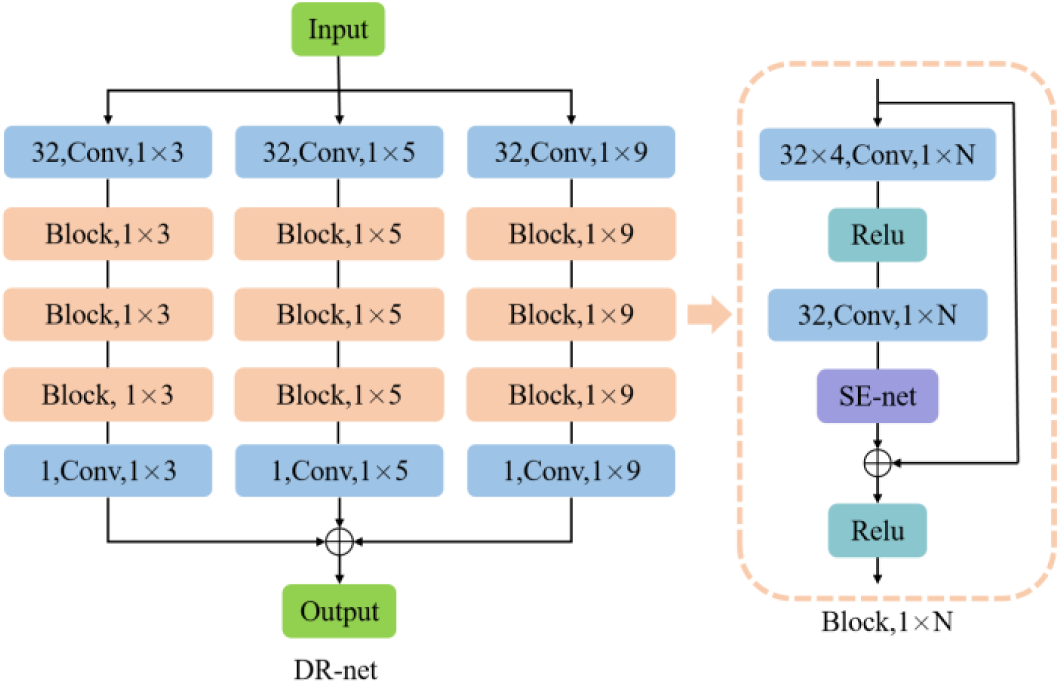
DR-net model structure.

**Fig. 5.**
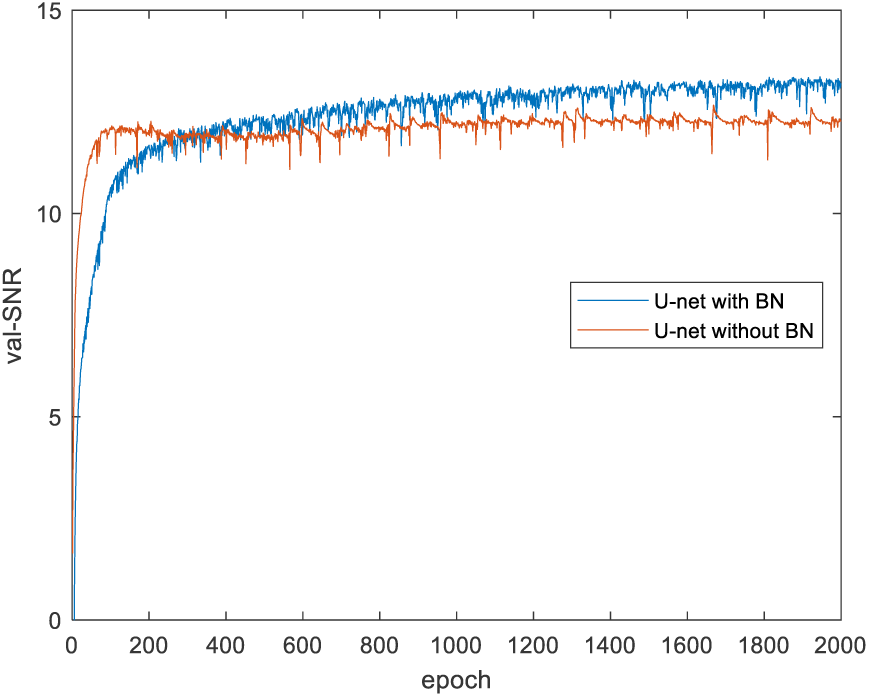
U-net with BN and U-net without BN. Comparison on Group1 validation set

**Fig. 6.**
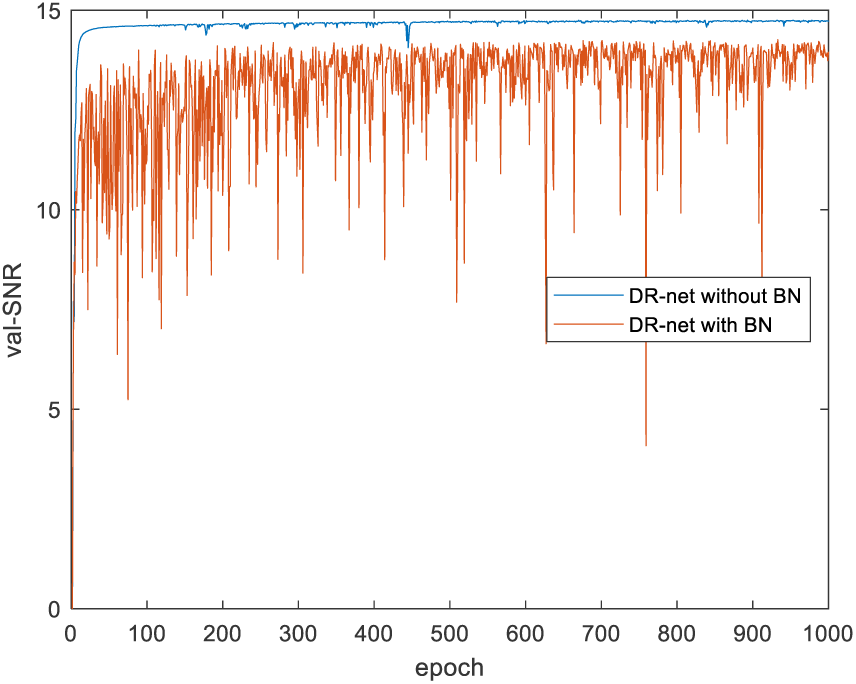
DR-net with BN and DR-net without BN. Comparison on Group1 validation set.

**Fig. 7.**
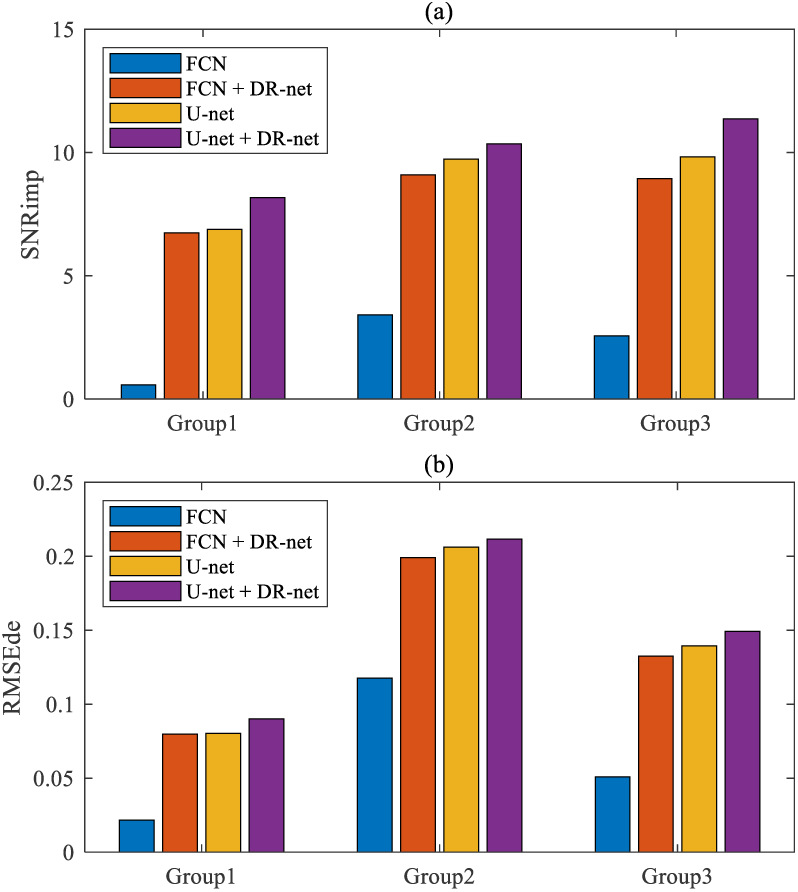
(a) *SNR*_*imp*_ and (b) *RMSE*_*de*_ of four ECG denoising methods tested on Group1, Group2, and Group3 test sets.

For this reason, we specially designed a multi-scale network for detailed restoration in phase two (as shown in Fig. 4), and we named it DR-net. To enhance DR-net’s ability to recover lost detail waveforms, the network design has two inputs: one input is the output of U-net after the first stage, and the other is the original noisy signal. The output is the corresponding ground-truth ECG. Adding the noise-convolved ECG to the input is helpful for DR-net to learn the missing part in the first stage, and it is helpful for it to find the regular pattern of the corresponding waveform after the first stage. In order to prevent the loss of details due to resampling, DR-net does not include downsampling and upsampling but uses a multi-scale network. The structure of the network is divided into three branches, and convolution kernels of different sizes are used in the three branches, which are denoted as 3, 5, and 9. In each branch, a convolution (conv) is passed first, then three blocks, and then another conv. There are 2 convs and 1 SENet[26] module in the block. The first conv module expands the number of convolution kernels by four times to 4 * 32. The expansion of the convolution kernel is conducive to increasing the network’s ability to process high-dimensional information. SENet added to each block is conducive to the network’s learning of detailed features. In U-net and DR-net, the training method is the adam algorithm, and the parameter settings are the same, where lr = 0.001, beta_1 = 0.9, beta_2 = 0.999, epsilon = 1e-08, and clipvalue = 0.5.

In DR-net, the size of the convolution kernel is smaller than U-net. This setting is good for observing the details in a waveform. Strictly speaking, the second stage does not learn how to remove noise but rather learns to recover the effective part based on the first stage U-net denoising. If DR-net noise reduction is used in the first stage, or U-net is used in the second stage, the effect is poor. The order in which the two networks are used should not be changed, and the two networks work best together.

It should be noted here that BN is used in the U-net network, but it is not used in the one-dimensional DR-net network. In the improved one-dimensional U-net network, there are processes of downsampling and upsampling. Adding the BN layer can avoid the disappearance of the gradient and enhance the performance of the network. But in a one-dimensional DR-net network, multi-scale convolution is used instead of the downsampling and upsampling layers. If BN is added, the network will lose the flexibility of the scope, and eventually the network’s denoising performance will be reduced.

## V. Results

Taking Group1 as example, we compared the performance of U-net and DR-net before and after adding BN. It can be seen in Fig. 5 that when BN is not added to U-net, U-net basically enters a stable state after 500 epochs of training. It is difficult to continue to improve the SNR and eventually reach 12.66 db. When BN is added to U-net, the network’s performance gradually improves until it stabilizes at 1,500 epochs and eventually reaches 13.36 db. It can be seen in Fig. 6 that when BN is added to each convolutional layer in DR-net, it causes instability during training, and the SNR curve of the validation set oscillates in a wide range. After 1,000 epochs, the SNR on the final verification set reaches 14.26 db. Without BN, the SNR curve of the verification set converges fast and is relatively stable. It enters the convergence state within 100 epochs, and the SNR on the final verification set reaches 14.75 db.

Next, we compared the two-stage method proposed in this paper (U-net + DR-net) with the FCN, FCN+DR-net, and U-net methods. The structure and training method of FCN was derived from [17]. The obvious feature in FCN is to realize the encoding function by setting the stride of the convolution and to realize the decoding function by the deconvolutional layers. For a fair comparison, the number of layers and the size of the convolution kernel of the FCN network were set to be the same as those in U-net, and the network was also trained by 2,000 epochs. Result appears in the following were taken from the average result of 138 sets of test data, and the test data were not trained.

Group1 is an MA-dominant data set. For the noise-convolved ECG in the test set, average *SNR* = 5.19 db, average *RMSE* = 14.82 × 10^-2^; after FCN denoising, *SNR*_*imp*_ = 0.58 db, *RMSE*_*de*_ = 2.66 × 10^-2^; after FCN + DR-net denoising, *SNR*_*imp*_ = 6.74 db, *RMSE*_*de*_ = 7.99 × 10^-2^; after U-net denoising, *SNR*_*imp*_ = 6.88 db, *RMSE*_*de*_ = 8.04 × 10^-2^; after U-net + DR-net denoising, *SNR*_*imp*_ = 8.18 db, *RMSE*_*de*_ = 9.01 × 10^-2^.

Group2 is an EM-dominant data set. For the noise-convolved ECG in the test set, average *SNR* = 0.60 db, average *RMSE* = 29.20 × 10^-2^; after FCN denoising, *SNR*_*imp*_ = 3.41 db, *RMSE*_*de*_ = 11.77 × 10^-2^; after FCN + DR-net denoising, *SNR*_*imp*_ = 9.09 db, *RMSE*_*de*_ = 19.91 × 10^-2^; after U-net denoising, *SNR*_*imp*_ = 9.73 db, *RMSE*_*de*_ = 20.63 × 10^-2^; after U-net + DR-net denoising, *SNR*_*imp*_ = 10.35 db, *RMSE*_*de*_ = 21.16 × 10^-2^.

Group3 is a BW-dominant data set. For the noise-convolved ECG in the test set, average *SNR* = 2.78 db, average *RMSE* = 20.39 × 10^-2^; after FCN denoising, *SNR*_*imp*_ = 2.56 db, *RMSE*_*de*_ = 5.09 × 10^-2^; after FCN + DR-net denoising, *SNR*_*imp*_ = 8.95 db, *RMSE*_*de*_ = 13.25 × 10^-2^; after U-net denoising, *SNR*_*imp*_ = 9.82 db, *RMSE*_*de*_ = 13.95 × 10^-2^; after U-net + DR-net denoising, *SNR*_*imp*_ = 11.37 db, *RMSE*_*de*_ = 14.93 × 10^-2^.

Figure 7 shows the values of *SNR*_*imp*_ and *RMSE*_*de*_ when using FCN, FCN + DR-net, U-net, and U-net + DR-net in Group1, Group2, and Group3. The results show that the performance of U-net + DR-net is superior. Furthermore, the denoising result using DR-net + FCN is better than using FCN alone; the results of both *SNR*_*imp*_ and *RMSE*_*de*_ are greatly improved, which proves that the second stage DR-net has excellent waveform recovery function.

Table 1 displays the correlation coefficients (in the form of mean ± standard deviation) between the QRS and T waves of denoised ECG obtained by the four methods and the QRS and T waves of the ground-truth ECG in the three groups of experiments. The larger the mean value of P and the smaller the standard deviation, the higher the correlation between denoised ECG and ground-truth ECG, and the more stable the result. Except for the standard deviation of the QRS waveform correlation coefficient in Group2, the rest of the indicators of U-net + DR-net are the best results among the four methods. Whether FCN or U-net is set as the first stage model, adding DR-net in the second stage can improve the final denoising performance in most cases. In particular, the combination of FCN and DR-net can greatly improve the denoising effect over using FCN alone.

**TABLE I:** Correlation Coefficients Between QRS and T Wavesw of Denoised ECG and Ground-truth ECG

| Model | Group1 (MA-dominant) |  | Group2 (EM-dominant) |  | Group3 (BW-dominant) |  |
| --- | --- | --- | --- | --- | --- | --- |
|  | QRS | ST-T | QRS | ST-T | QRS | ST-T |
| FCN | 0.7480<br>$\pm 0.3468$ | 0.5805<br>$\pm 0.4142$ | 0.7164<br>$\pm 0.3887$ | 0.4667<br>$\pm 0.5138$ | 0.8245<br>$\pm 0.2976$ | 0.5793<br>$\pm 0.4375$ |
| FCN+DR-net | 0.9675<br>$\pm 0.04965$ | 0.8329<br>$\pm 0.2531$ | 0.9439<br>$\pm 0.1122$ | 0.7374<br>$\pm 0.3315$ | 0.9668<br>$\pm 0.0592$ | 0.8233<br>$\pm 0.2735$ |
| U-net | 0.9808<br>$\pm 0.0270$ | 0.8671<br>$\pm 0.2551$ | 0.9643<br>$\pm 0.1006$ | 0.7724<br>$\pm 0.3335$ | 0.9834<br>$\pm 0.0211$ | 0.8550<br>$\pm 0.2587$ |
| U-net+DR-net | 0.9916<br>$\pm 0.0252$ | 0.8809<br>$\pm 0.2542$ | 0.9745<br>$\pm 0.1044$ | 0.7962<br>$\pm 0.3325$ | 0.9940<br>$\pm 0.0207$ | 0.8788<br>$\pm 0.2579$ |

During the experiment, we found that FCN could not achieve the denoising effect described in the paper. The author used the data from MIT-BIH during the experiment and extracted 200 fragments from each set of data. Our analysis is that after the disruption, the training set, the verification set, and the test set were all leaked.

Figure 8 shows a segment of ECG signal with atrial premature beats. This segment of the signal is characterized by a deep downward QRS complex. As can be seen in Fig. 8(a) and (b), the segment of noise-convolved ECG belongs to high-quality ECG with an *SNR* of 15.86 db. As can be seen in Fig. 8(c) and (e), when FCN or U-net is used for the first stage of denoising, the downward spikes and peaks will be suppressed to a certain extent. At this time, the FCN method’s *SNR*_*imp*_ = −6.90 db, and the U-net method’s *SNR*_*imp*_ = −1.38 db, the two methods of the first stage reduce the signal quality of the signal in this segment, but the degree of decline is different. However, it can be seen from Fig. 8(d) and (f) that by adding the second stage with DR-net, the suppressed downward spikes and T waves are effectively restored. At this time, the FCN + DR-net method’s *SNR*_*imp*_ = 3.44 db, and the U-net + DR-net method’s *SNR*_*imp*_ = 5.35 db, indicating that the DR-net in the second stage plays an important role in waveform recovery.

**Fig. 8.**
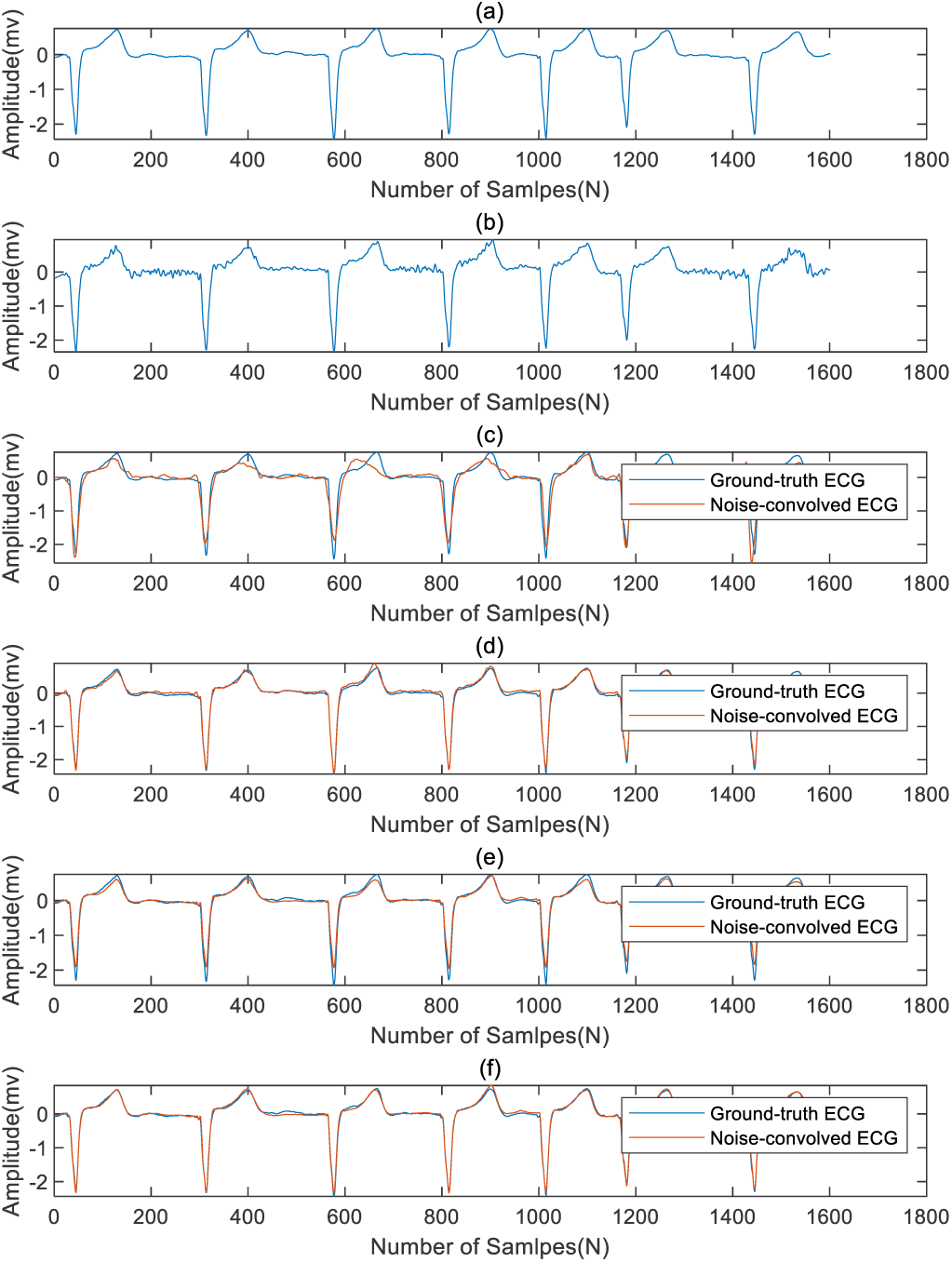
(a) Ground-truth ECG. (b) Noise-convolved ECG. (c) Denoised ECG by FCN. (d) Denoised ECG by FCN + DR-net. (e) Denoised ECG by U-net. (f) Denoised ECG by U-net + DR-net.

Figure 9 shows a segment of an ECG signal from Group1 with a signal-to-noise ratio of 5.42 db. The characteristics of this signal are the round bluntness of the R wave and the downward movement of the ST segment. As can be seen in Fig. 9(b), this signal is mainly affected by EMG. It can be seen in Fig. 9(c) and (d) that the denoising effect of FCN + DR-net is improved compared with the denoising effect of FCN, but it is still unsatisfactory in the recovery of the ST-T segment since the downward shift of the ST segment is not reflected in the denoised waveform. But no such problem occurs in U-net. From Fig. 9(e), it can be seen that, despite the flaws in the amplitude of the peak, the basic characteristics of the waveform are relatively complete. As shown in Fig. 9(f), the amplitude of the peak is restored after the addition of DR-net.

**Fig. 9.**
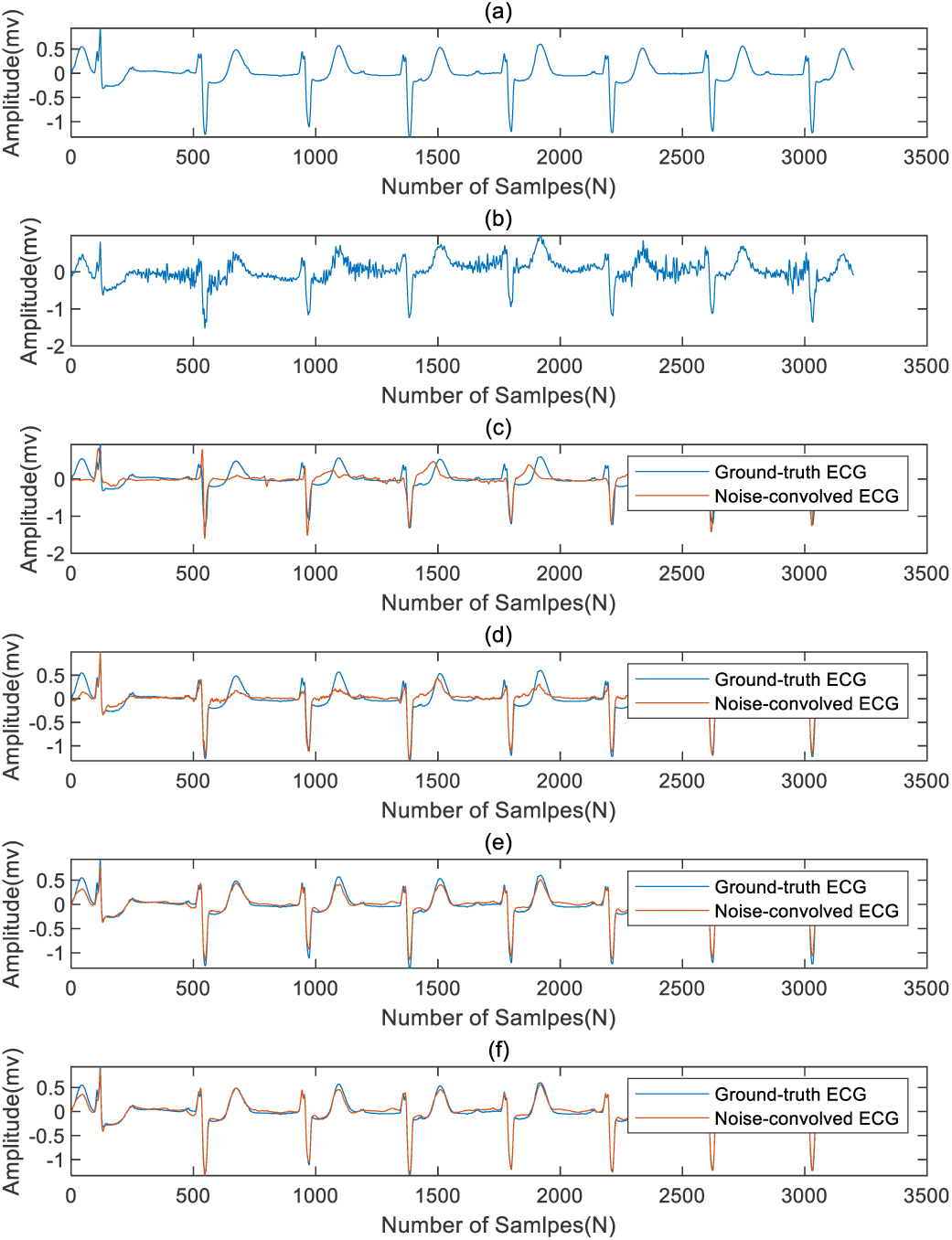
(a) Ground-truth ECG. (b) Noise-convolved ECG. (c) Denoised ECG by FCN. (d) Denoised ECG by FCN + DR-net. (e) Denoised ECG by U-net. (f) Denoised ECG by U-net + DR-net.

Figure 10 shows a segment of an ECG signal from Group2 with *SNR* = 2.36 db. The ECG signal in this segment is mainly affected by EM, which causes a large degree of variation in the T wave of the ECG. The R-wave position also shifts. By analyzing Fig. 10, it is possible to draw a similar conclusion to that in Fig. 9. U-net has achieved better results than the FCN method, and DR-net must be added to further improve the signal quality of denoised ECG.

**Fig. 10.**
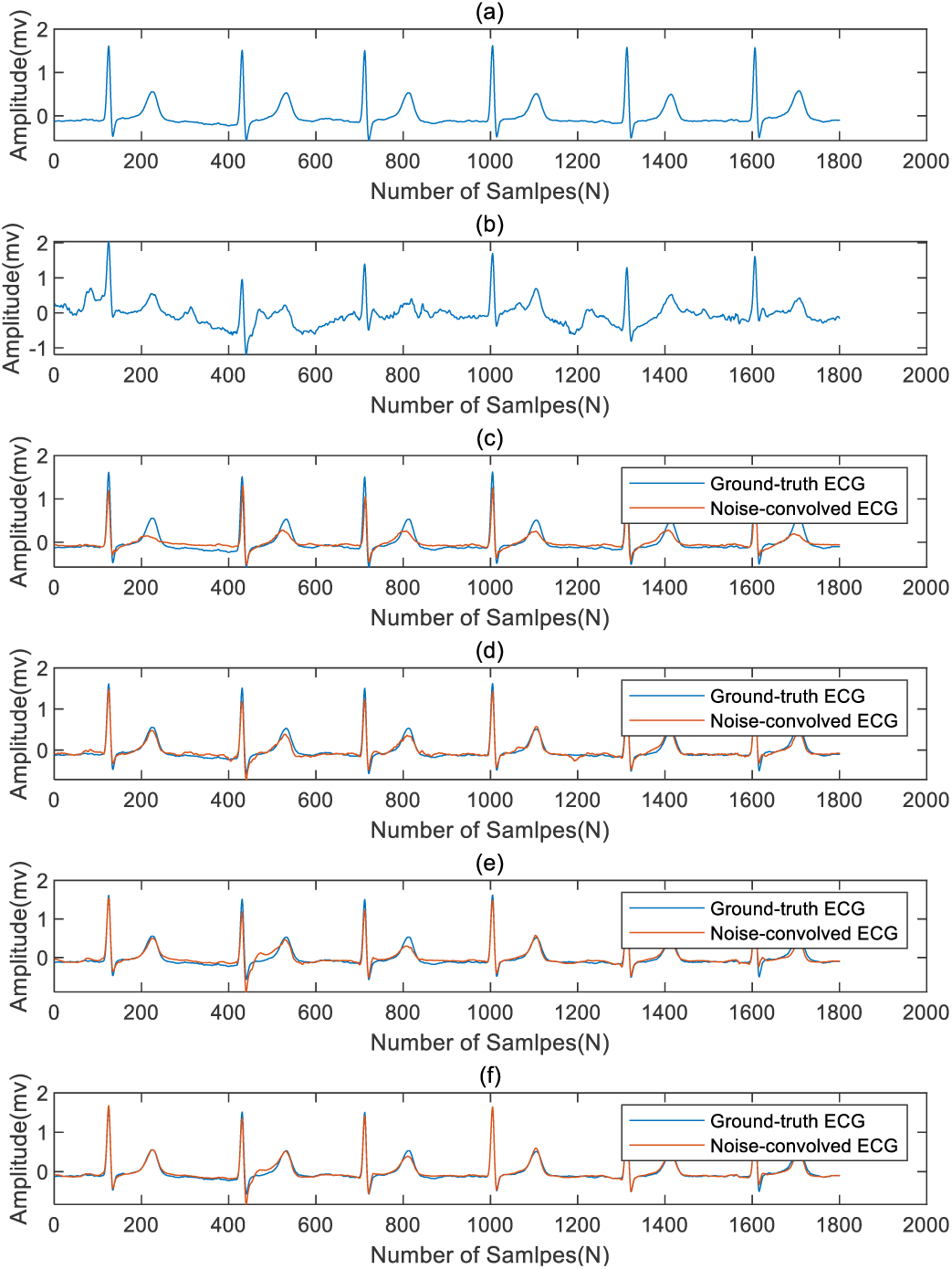
(a) Ground-truth ECG. (b) Noise-convolved ECG. (c) Denoised ECG by FCN. (d) Denoised ECG by FCN +DR-net. (e) Denoised ECG by U-net. (f) Denoised ECG by U-net +DR-net.

Figure 11 shows a segment of an ECG signal from Group3 with *SNR* = −3.1343 db. The signal in this segment is mainly affected by BW. The characteristic of this segment is that the third heartbeat is an inserted ventricular premature beat. The heartbeat shows a wide QRS complex and a large T wave inversion. Comparing Fig. 11(c) and (d) in terms of the difference in inserted ventricular premature beats, FCN + DR-net recovers the wide QRS complex better than FCN alone. However, the shape of the inverted T wave is not restored. Comparing Fig. 11(e) and (f), it can be found that both U-net and U-net + DR-net methods can achieve good results: U-net *SNR*_*imp*_ = 17.29, and U-net + DR-net *SNR*_*imp*_ = 18.55, which are basically at the same level.

**Fig. 11.**
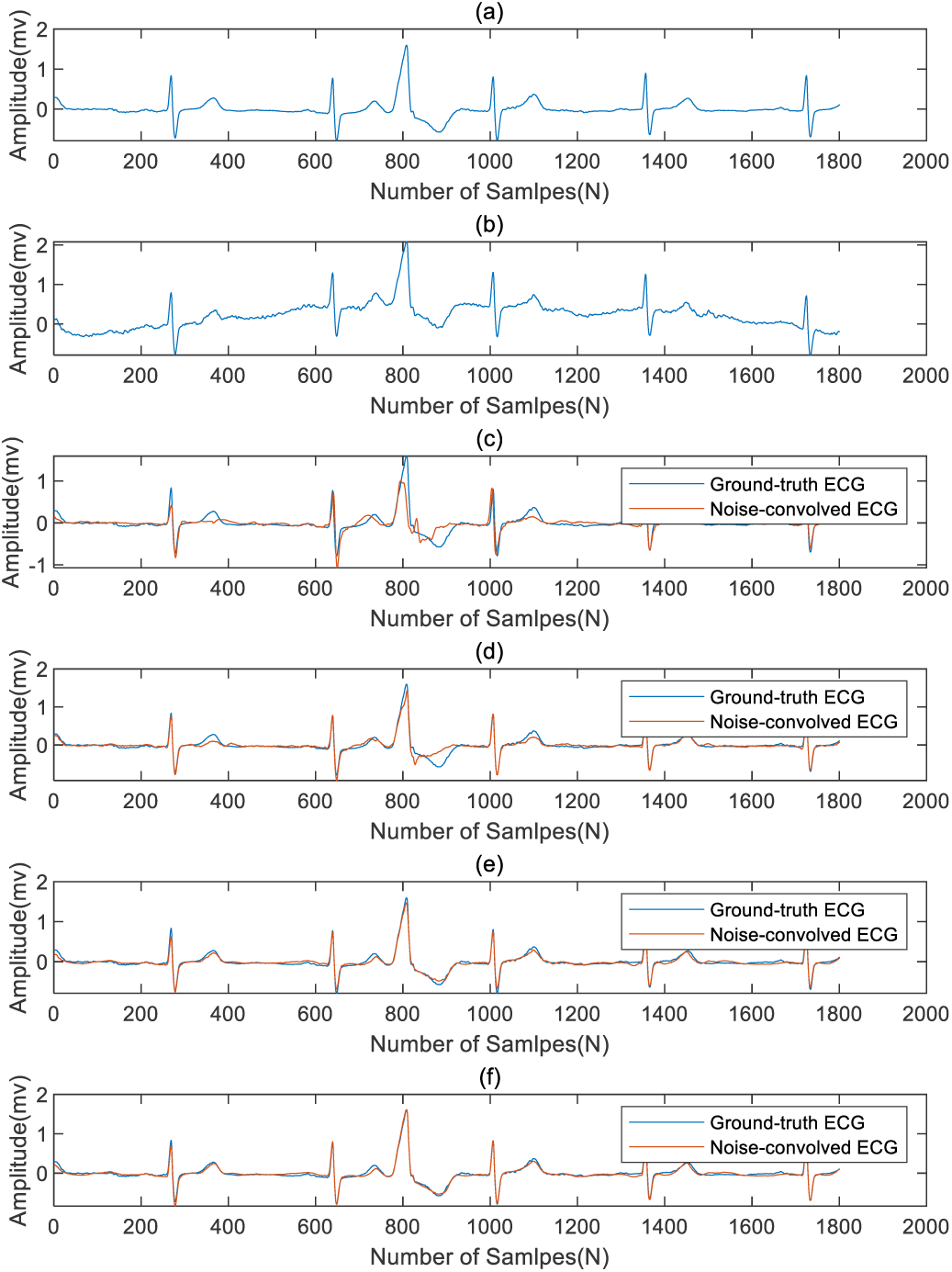
(a) Ground-truth ECG. (b) Noise-convolved ECG. (c) Denoised ECG by FCN. (d) Denoised ECG by FCN +DR-net. (e) Denoised ECG by U-net. (f) Denoised ECG by U-net +DR-net.

With reference to Figs. 9 and 11, it can also be explained that when the waveform of the first stage of denoising causes a large distortion, it is difficult for DR-net to recover in the second stage. Therefore, we think that using U-net in the first stage and DR-net in the second stage can achieve the best results.

## VI. Summary

This paper presents a novel two-stage denoising method for removing noise from ECG signals that are contaminated by baseline drift, muscle artifacts, and electrode motion. We propose an improved one-dimensional U-net, which improves the size of the convolution kernel and the structure of the network so that it can better perform the task of denoising. We specially designed DR-net for detailed restoration in the second stage. The network can continue to improve the signal quality based on the first stage and can effectively reduce the error between the denoised signal and the characteristic waveform of the real signal. In ECG signals with a very high signal-to-noise ratio, U-net or FCN will cause signal quality degradation, but after the second stage with DR-net, the lost information can be recovered to a certain extent. Denoising and the preservation of effective details are somewhat contradictory, but the two-stage method proposed in this paper can achieve both the elimination of noise and the preservation of effective details to a large extent. We believe that the proposed method has good application prospects in clinical practice.

The network proposed in this paper was developed using Python, using Keras for simple prototyping, and using TensorFlow as a back-end deep learning library. The workstation specifications for training the model include an NVIDIA GPU: Tesla K40m and an 11GB memory. U-net was trained for 2,000 epochs, and DR-net was trained for 1,000 epochs.

